# Multi-locus and long amplicon sequencing approach to study microbial diversity at species level using the MinION™ portable nanopore sequencer

**DOI:** 10.1101/117143

**Authors:** Alfonso Benítez-Páez, Yolanda Sanz

**Affiliations:** Microbial Ecology, Nutrition & Health Research Unit. Institute of Agrochemistry and Food Technology Institute (IATA-CSIC), Valencia, Spain.

**Keywords:** MinION, Nanopore sequencer, Ribosomal operon, Long amplicon sequencing, Microbial diversity, Long-read sequencing

## Abstract

**Background:** The miniaturised and portable DNA sequencer MinION™ has demonstrated great potential in different analyses such as genome-wide sequencing, pathogen outbreak detection and surveillance, human genome variability, and microbial diversity. In this study, we tested the ability of the MinION™ platform to perform long amplicon sequencing in order to design new approaches to study microbial diversity using a multi-locus approach.

**Results:** After compiling a robust database by parsing and extracting the *rrn* bacterial region from more than 67,000 complete or draft bacterial genomes, we demonstrated that the data obtained during sequencing of the long amplicon in the MinION™ device using R9 and R9.4 chemistries was sufficient to study two mock microbial communities in a multiplex manner and to almost completely reconstruct the microbial diversity contained in the HM782D and D6305 mock communities.

**Conclusions:** Although nanopore-based sequencing produces reads with lower per-base accuracy compared with other platforms, we presented a novel approach consisting of multi-locus and long amplicon sequencing using the MinION™ MkIb DNA sequencer and R9 and R9.4 chemistries that help to overcome the main disadvantage of this portable sequencing platform. Furthermore, the nanopore sequencing library constructed with the last releases of pore chemistry (R9.4) and sequencing kit (SQK-LSK108) permitted to retrieve the higher level of 1D read accuracy sufficient to characterize the microbial species present in each mock community analysed. Improvements in nanopore chemistry, such as minimising base-calling errors and new library protocols able to produce rapid 1D libraries, will provide more reliable information in near future. Such data will be useful for more comprehensive and faster specific detection of microbial species and strains in complex ecosystems.

## Background

During the last two years, DNA sequencing based on single-molecule technology has completely changed the perception of genomics for scientists working in a wide range of scientific fields. This new perspective is not only supported by the technology itself but also by the affordability of these sequencing instruments. In fact, unprecedentedly, Oxford Nanopore Technologies (ONT) released the first miniaturised and portable DNA sequencer in early 2014, within the framework of the MinION™ Access Programme. Recently, the MARC consortium (MinION Analysis and Reference Consortium) has published results related to the study of the reproducibility and global performance of the MinION™ platform. These results indicate that this platform is susceptible of a large stochastic variation, essentially derived from the wet-lab and MinION™ operative methods, but also that variability has minimal impact on data quality [1].

The coordinated and collaborative work and mutual feedback between industry and the scientific community have enabled ONT to develop rapidly towards improving its portable platform for DNA sequencing, minimizing the stochastic variation during DNA library preparation. Consequently, in late Autumn 2015, ONT released MkIb, the latest version of MinION™, and in April 2016 the fast mode chemistry (R9) was released, increasing the rate of sensing DNA strands from 30-70 to 280-500 bp/sec and reaching up to 95% of per-base accuracy in 2D reads (Clive G. Brown, CTO ONT, personal communication).

One of the most attractive capabilities of the MinION™ platform is the sequencing and assembly of complete bacterial genomes using exclusively nanopore reads [2] or through hybrid approaches [3, 4]. Notwithstanding, the MinION™ platform has also been demonstrated useful in other relevant areas including: human genetic variant discovery [5, 6], detection of human pathogens [7, 8], detection of antibiotic resistance [9, 10], and microbial diversity [11, 12]. Regarding the latter, microbial diversity and taxonomic approaches are common and in high demand to analyse the microbiota associated to a wide variety of environment- and human-derived samples. However, these analyses are greatly limited by the short-read strategies commonly employed. Thanks to improvements in the chemistry of the most common, popular sequencing platforms in recent years, it is now possible to characterise microbial communities in detail, down to the family or even genus level, using genetic information derived from roughly 30% (~500nt) of the full 16S rRNA gene. Despite the massive coverage achieved with short-read methods, the limitation in terms of read length means taxonomic assignment at the species level is still unfeasible. For instance, taxonomy strategies based on short-reads from Illumina MiSeq platform offer a limited information that underestimates the microbial diversity of complex samples when compared with alternative approaches based on long DNA reads [13]. Consequently, implementation of long-read sequencing approaches to study larger fragments of marker genes will permit the design of new studies to provide evidence for the central role of precise bacterial species/strains in a great variety of microbial consortia. Recent studies at this regard have showed important advances in taxonomy analysis using long reads generated by single molecule technologies [11, 14, 15], indicating that the expansion or inclusion of more hypervariable regions in the analysis overcomes the disadvantage of working with error-prone DNA reads. With respect to the above, we have recently explored the performance of the MinION™ device. Our study demonstrates that data obtained from sequencing nearly full-length 16S rRNA gene amplicons is feasible to study microbial communities through nanopore technology [11]. We wanted to move a step forward in this type of strategy, thus gaining more specificity when including several hypervariable markers in the analysis, at sequence and structural level, by designing a multi-locus and long amplicon sequencing method to study microbial diversity. At the same time, we also wanted to explore the affordability of the MinION™ technology to perform microbial diversity analyses by multiplexing several samples in one single MinION™ flowcell. Accordingly, here we present a study of the 16S, 23S, and the internal transcribed spacer (ITS, that frequently encodes tRNA genes) simultaneous sequencing, using the MinION™ MkIb device and R9 chemistry, with prior generation of ~4.5kb DNA fragments by amplifying the nearly full-length operon encoding the two larger ribosomal RNA genes in bacteria, the *rrn* region (*rrn* hereinafter). We have studied the *rrn* of two mock microbial communities, composed of genomic DNA from 20 and 8 different bacterial species, obtained respectively from BEI Resources and ZYMO Research Corp., using the MinION™ sequencing platform in multiplex configuration.

## Data description

The R9 raw data collected in this experiment was obtained as fast5 files using MinKNOW™ v0.51.3.40 (Oxford Nanopore Technologies) after conversion of electric signals into base calls via the Metrichor™ agent v2.40.17 and the Barcoding plus 2D Basecalling RNN for SQK-NSK007 workflow v1.107; whereas the R9.4 raw data was generated by MinKNOW™ v1.5.5 (Oxford Nanopore Technologies) with the respective local basecalling algorithm implemented in that version of the MinION™ controller software. Base-called data passing quality control and filtering were downloaded and data was converted to fasta format using the *poRe* package [16]. Fast5 raw data can be accessed at the European Nucleotide Archive (ENA) under the project ID PRJEB15264. Only two data sets were generated after a sequencing run of MinION™ MkIb.

## Analysis

### Defining the arrangement of the rrn region

The complete or partial gene sequence of the RNA attached to the small subunit of the ribosome is classically used to perform taxonomy and diversity analysis in complex samples containing hundreds of microbial species. In the case of bacterial species, the 16S rRNA gene is the most widely used DNA marker for taxonomic identification of a particular species, given the relatively high number of hypervariable regions (V1 to V9) present across its sequence. Nowadays, it is possible to study the complete or almost full-length sequence of the 16S rRNA molecule thanks to single-molecule sequencing approaches [11, 14, 15, 17]. The identification of complex microbial communities at species-level with raw data obtained from MinION™ or PacBIO platforms is improving; however, uncertainty in taxonomic assignation is still noteworthy given the high proportion of errors in their reads. While future technical advances may improve the quality of DNA reads generated by third generation sequencing devices, new strategies can also be adopted to enhance the performance of these approaches. Consequently, we postulate that a good example of this is to study a common multi-locus region of the bacterial genome, which enables the simultaneous study of more variable regions and locus arrangements (sequence and structural variation), such as the operon encoding the ribosomal RNA. Using a complex sample where hundreds of microbial species are potentially present (DNA from human faeces) we carried out preliminary experiments to amplify the *rrn*. We observed that from the hypothetical configurations envisaged for the *rrn* (Figure 1A), we only obtained a clear amplification using the primer pairs S-D-Bact-0008-c-S-20 and 23S-2241R, indicating that the *rrn* preferentially seems to be transcriptionally arranged as follows: 16S-ITS-23S. A detailed evaluation of the fragment size determined that main PCR products ranged from 4.3 to 5.4kbp (Figure 1B-D), being consistent with the expected size of PCR products amplifying the 16S, ITS, and 23S regions from several microbial species. The next step involved designing a multiplex sequencing approach to try to analyse more than one sample per sequencing run in one flowcell of MinION™; therefore, the primers were re-designed to include a distinctive barcode region at 5′ (Table 1). During PCR of the *rrn* we tagged the amplicon derived from the mock community HM782D with the barcode *bc01* in a dual manner, whereas the amplicons derived from sample D6305 were tagged with barcode *bc08* in similar way. Parallel experiments were conducted on HM782D and D6305 DNA, with comparative aims, using a conventional protocol of microbial diversity analysis and consisting of the V4-V5 16S amplicon sequencing by Illumina MiSeq paired-end approach (see methods).

**Figure 1.**
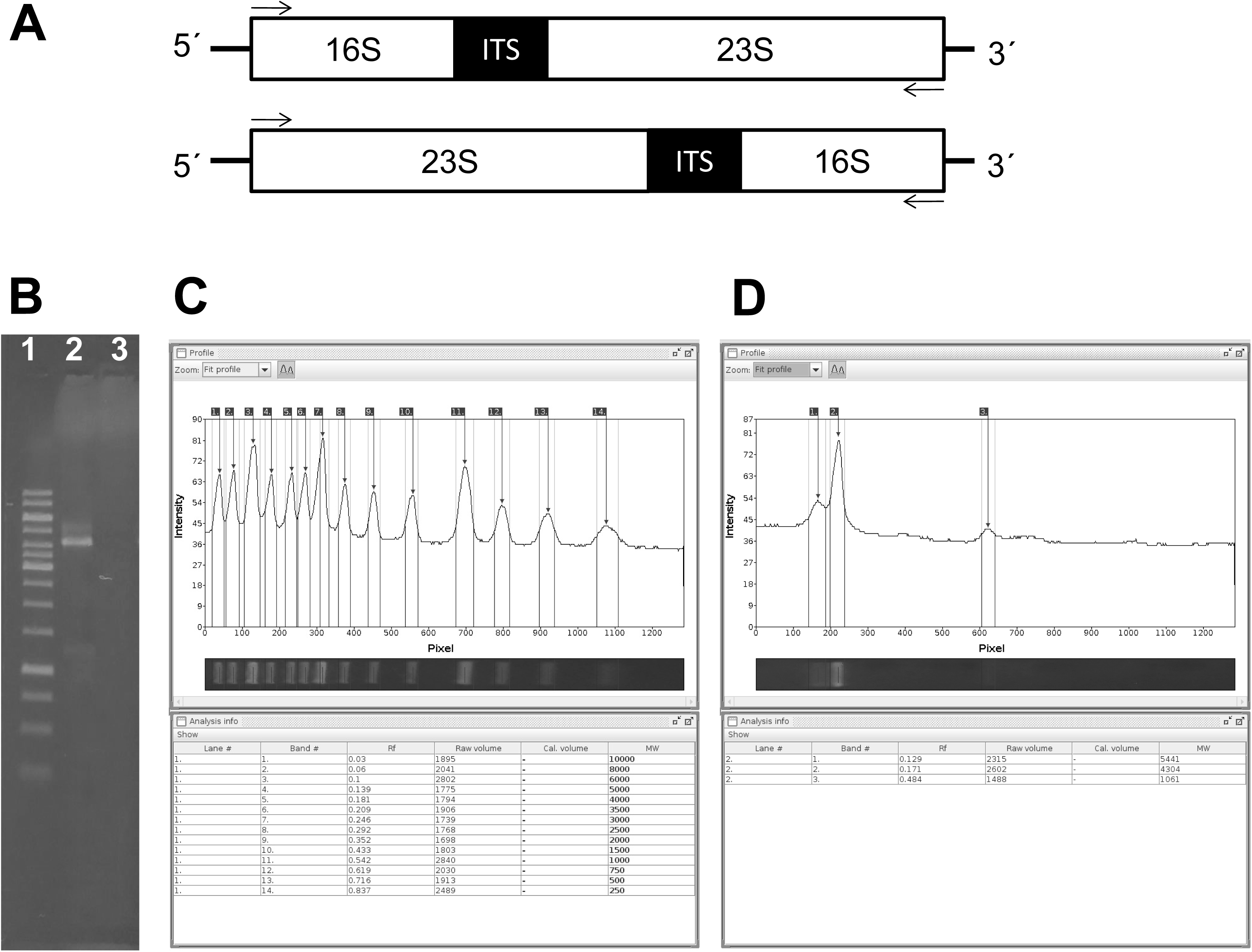
Organization of the *rrn* region in bacteria. A - hypothetical transcriptional arrangements expected for *rrn* and tested experimentally using two sets of primer pairs (see small arrows drawn in each configuration). B - Agarose gel electrophoresis of PCR reactions performed under the two hypothetical arrangements of *rrn*; lanes: 1) 1kb ruler (Fermentas), 2) PCR reaction from the top configuration in panel A, 3) PCR reaction from the bottom configuration in panel A. The GelAnalyser Java application was used to perform the band size analysis of the 1kb ruler standard (C) and the amplicons obtained from human faecal DNA (D).

**Table 1.**
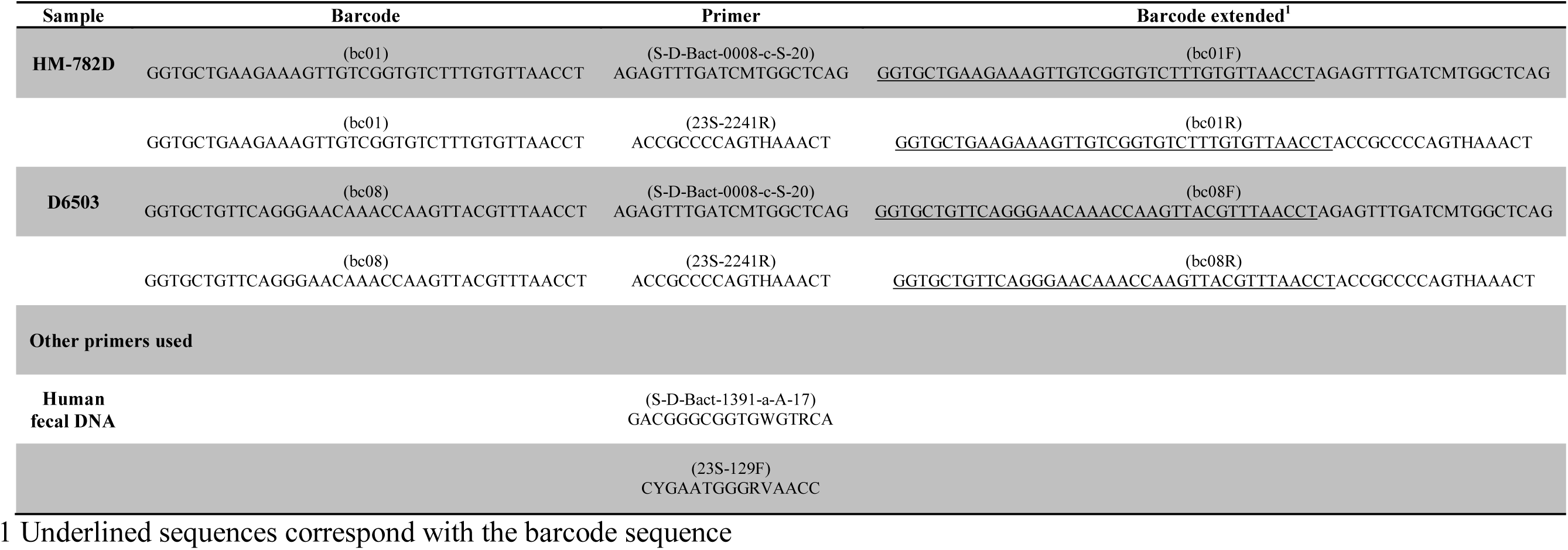
Barcodes and primers used to generate amplicon libraries.

### The reference database

One of the major handicaps when proposing this new *rrn* region to be used for taxonomy analysis is the need to compile a reference database to compare the reads produced by MinION™ device. Therefore, we proceeded to parse the genetic information of over 67,000 bacterial genomes whose sequences are publicly available in GenBank at NCBI. In this way, we retrieve and compile more than 47,000 *rrn* sequences that were subject of a clustering analysis to reduce the level of redundancy and to disclose the variability intrinsically associated to the *rrn* itself and to its individual components as well (Figure 2).

**Figure 2.**
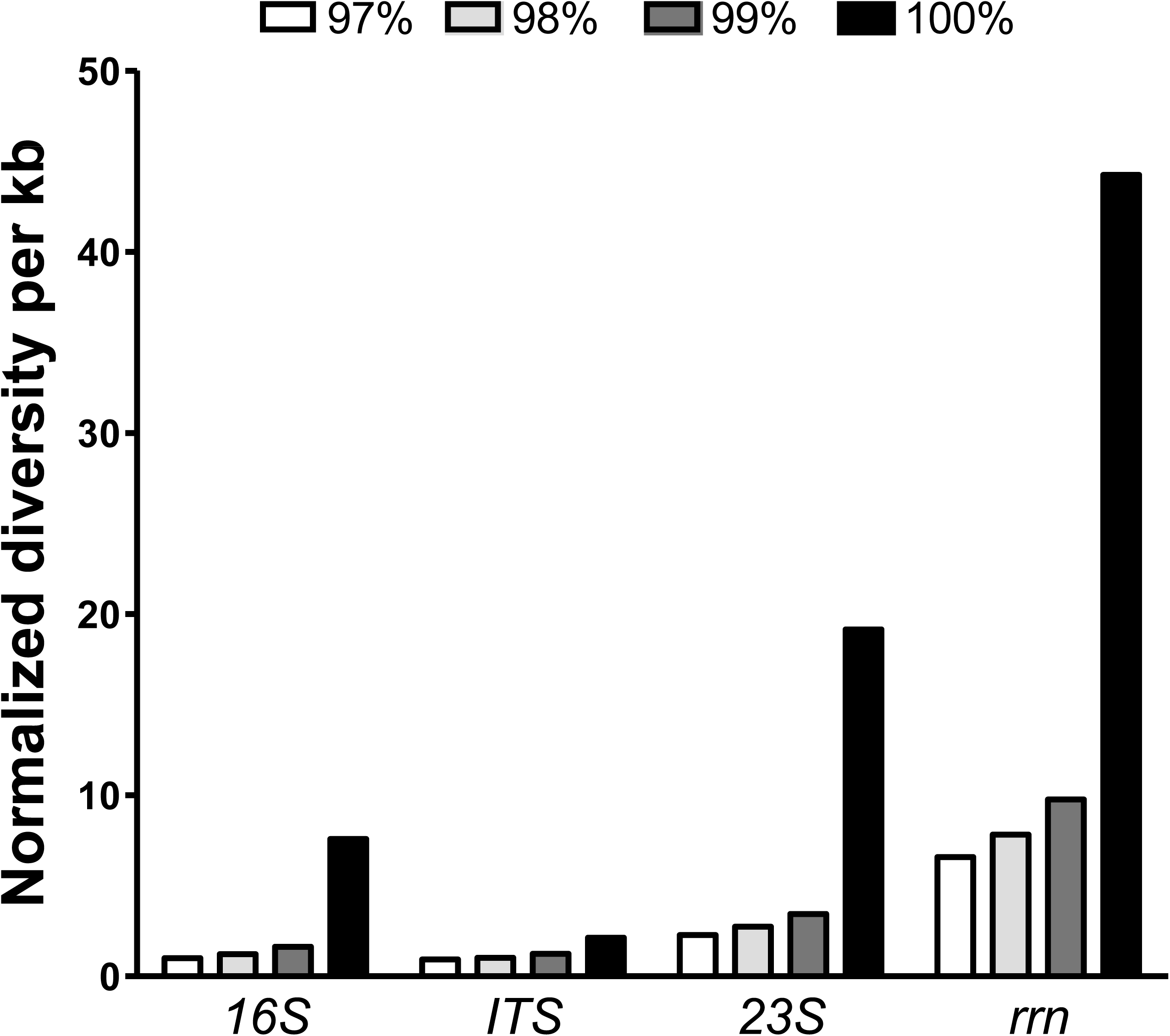
Variability of the *rrn* region and its functional domains. The *rrn* database compiled after parsing more than 67,000 draft and complete bacterial genomes was assessed by clustering analysis at different levels of sequence identity: 97 (white bars), 98 (light grey bars), 99 (dark grey bars), and 100% (black bars). For comparative aims, the functional DNA sequences encoded into the *rrn* region were also individually studied. The normalized diversity (y axis) resulted from calculate the number of clusters obtained for each analysis normalized with the median sizes of respective regions in terms of kb, and referenced against the value obtained for 16S sequences clustered at 97%, the canonical threshold for species assignment.

After normalization of cluster numbers against the median size of respective regions analyzed and referenced against the numbers obtained for 16S region at 97% sequence identity, we found that *rrn* region comprising the 16S, ITS, and 23S coding regions exhibits more than 4-fold more variation than that observed for the 16S molecule alone (at 100% sequence identity). As expected, the 23S region exhibited more diversity by containing more hypervariable regions than 16S region and getting almost 2-fold more diversity. Strikingly, the ITS regions showed similar levels of genetic diversity despite to have almost one fourth of the size of 16S region in average. When parsing the genetic information of over 67,000 bacterial genomes, we observed the ITS region frequently encodes one or several tRNA genes and it possess a high variability in terms of length as well. Consequently, the variability observed in the *rrn* was the largest observed and thought to be meaningful for the aims of this study. We obtaining data supporting the above notion by searching the number of *rrn* clusters (at 100% identity) matching with the most predominant species in the database, thus retrieving 1,713, 1,276, and 1,273 *rrn* clusters annotated for *Escherichia coli*, *Streptococcus pneumoniae*, and *Staphylococcus aureus*, respectively. In consequence, the *rrn* is able to accumulate enough sequence variability to discern taxonomy even at strain level.

### Performance of the R9 chemistry

Once we could compile a reference database for comparison aims, we proceeded with the amplicon library construction and sequencing run obtaining raw data consisting of 17,038 reads and almost all were classified as 1D reads. For general knowledge, the DNA reads derived from the MinION™ device can be classified into three types: ‘1D template’, ‘1D complement’, and ‘2D’ reads. The latter, 2D reads, are products of aligning and merging sequences from the template (read from leader adapter) and complement reads (a second adapter called hairpin or HP adapter must be generated), produced from the same DNA fragment. These contain a lower error rate, owing to strand comparison and mismatch correction. In addition to the technical issues indicative of a bad ligation of the HP adapter, we obtained 93% of reads (~15,900 reads) during the first 16h of run; thus, we obtained lower sequencing performance after reloading with the second aliquot of the sequencing library and extended the run for another 24h (40h in sum). The fasta sequences were filtered by retaining those between 1,500 and 7,000 nt in length, obtaining at least enough sequence information to compare a DNA sequence equivalent to the 16S rRNA gene length. After this filtering step, we retained 72% of sequences (12,278) and then we performed the respective barcode splitting. For this purpose, we modified the default parameters of the “split_barcodes.pl” perl script (Oxford Nanopore Technologies) by incorporating the information of the extended barcodes (Table 1), rather than the barcode information alone, and simultaneously increased the stringency parameter to 25 (14 by default). Afterwards the concatenation of reads were obtained from respective forward and reverse extended barcodes, then we retrieved a total of 2,019 (52% from forward and 48% from reverse barcodes) and 1,519 (53% from forward and 47% from reverse barcodes) 1D reads for HM782D and D6305 mock communities, respectively. Read-mapping was performed against the *rrn* database, compiling more than 22,000 *rrn* regions, retrieved from more than 67,000 genomes available in GenBank (see Availability of supporting data). The taxonomy associated to the best hit based on the competitive alignment score followed by filtering steps (see methods) was used to determine the structure of each mock community. The MinION™ sequencing data produced the microbial structure presented in Figure 3 for the mock communities HM782D and D6305, respectively.

**Figure 3.**
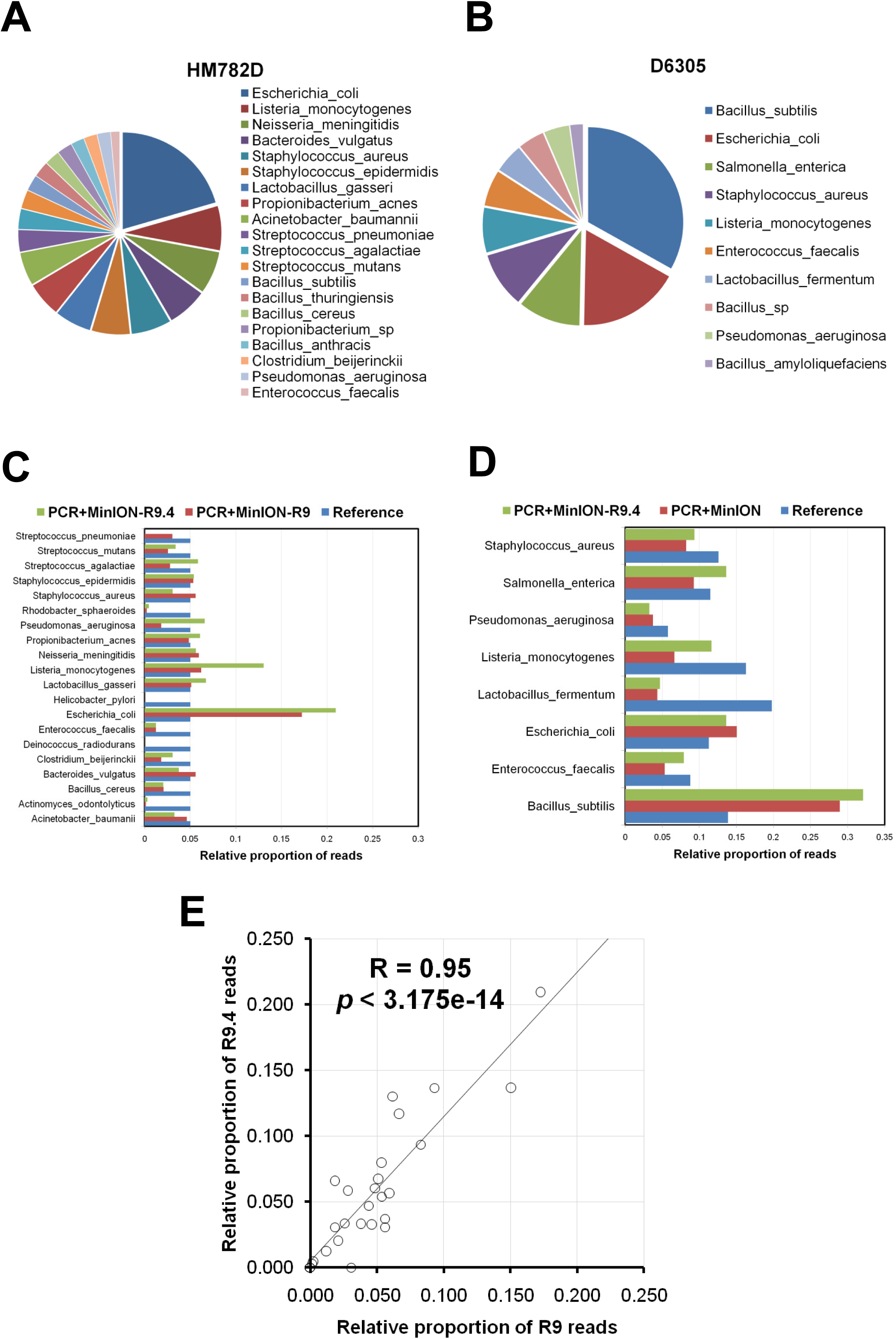
Microbial structure of the mock communities. A and B - microbial species and respective relative proportions determined to be present in the HM782D and D6305 mock communities, respectively, following the analysis of raw data obtained from *rrn* amplicon sequencing in the MinION™ and chemistry R9. C and D - Comparative analysis of the expected microbial species and proportions against the data obtained after mapping of reads generated by a 4.5kbp amplicon PCR and sequenced in MinION™ device with R9 and R9.4 chemistries, for HM782D and D6305 respectively. E - Linear correlation analysis of relative read proportions obtained for all bacterial species present in HM872D and D6305 mock communities with R9 and R9.4 chemistries.

Figure 3 shows the bacterial species and their respective relative proportions retrieved from the analysis of the mock communities HM782D and D6305, respectively. With respect to the HM782D mock community, we were able to recover 20 representative species, accounting for 16 out of 20 species present in that artificial community (Figure 3A). However, the remaining four species that apparently are absent in this community have a close relationship to others detected correctly, namely *Bacillus subtilis, Bacillus thuringensis, Bacillus anthracis*, and *Propionibacterium sp.* Furthermore, we were unable to report the presence of just four species present in HM782D because proportions of *Rhodobacter sphaeroides* and *Actinomyces odontolyticus* were below the predominance threshold (1%), being present in 0.25 and 0.12%, respectively. Similarly, other 40 different species but close to that present in the HM782D mock community (*Bacillus* spp., *Streptococcus* spp. *Clostriudium* spp., *Neisseria* spp., *Staphylococcus* spp, and *Listeria* spp.) had minor representation in data derived from *rrn* sequencing. With respect to *Rhodobacter sphaeroides* and *Actinomyces odontolyticus* lower proportions, we have previously demonstrated that the low levels of 16S reads are a consequence of amplification bias derived from the PCR reaction and not from sequencing itself [11]. In this case, the new primer pair used to generate the long amplicons would seem to work more efficiently than those previously used, but apparently they still present issues at bacterial coverage level. When we revised the whole taxonomy contained in our *rrn* database, the compiling of non *rrn* regions for *Deinococcus radiodurans* and *Helicobacter pylori* partially explained the lack of these species in HM782D analysed by the present approach. However, a new alignment process using individual 16S and 23S rRNA sequences obtained from GenBank and including those for *D. radiodurans* and *H. pylori*, respectively, demonstrated that at least *D. radiodurans* could be identified in a higher proportion than *A. odontolyticus* and *R. sphaeroides*, albeit in a lower proportion than our predominance threshold. Regarding the results obtained from the D6305 mock community, we found a total of 10 bacterial species present in this mixed DNA sample, eight of them matched the expected structure of the community, and additionally 18 close species had minor representation (*Bacillus* spp., *Enterococcus* spp., *Klebsiella* spp., *Lactobacillus* spp., *Streptococcus* spp., and *Staphylococcus* spp.). Using the MinION™ data we were able to recover 100% of the species present in this sample and the two additional members identified also have a close relationship within the *Bacillus* genus, as observed in the HM782D sample (Figure 3B). We have determined that coverage needed to retrieve all expected species in a non-even mock community with an abundance above 1% is ~13X in terms of the number of species of that community.

When compared to reference values and proportions theoretically expected for the species present in the two mock communities, we observed some deviations that were greater in certain species. Particularly, in the HM782D sample the lowest coverage biases were observed for *Actinomyces odontolyticus* (-5.36)*, Rhodobacter sphaeroides* (-4.36), and *Enterococcus faecalis* (-2.04). This indicates that such species, in addition to *D. radiodurans* and *H. pylori*, are more difficult to detect with the primers and PCR used here. By contrast*, Escherichia coli* (1.79) seems to be preferentially amplified, given that this species exhibited the highest positive coverage bias value (Figure 3C). We again found that coverage bias is linearly correlated with PCR products generated by quantifying *E.coli, L. gasseri*, and *B. vulgatus* amplicons (Pearson’s r = 0.82, p = 0.047), data indicating that there are not major issues during taxonomy assignation by over-representation of certain species in the reference database. The values obtained for D6305 were more homogeneous, and the lowest coverage bias was observed for *Lactobacillus fermentum* (-2.18) (Figure 3D). Additional analysis indicated that there was not significant correlation between coverage bias and GC content in *rrn.* Although the low coverage bias for some species can be solved by selecting another pair of primers, the ability to recover almost all of them, at least in a low proportion, in itself represents an important attribute of this approach for inter-sample comparisons. Interestingly, we observed a similar pattern of overrepresentation of *Bacillus* spp. sequences (>50%) in D6305 sample but not for *Escherichia* spp. sequences (~4%) in the HM782D mock community when Illumina MiSeq data was assessed (Figure 3C-D).

The high error rate of the 1D reads (ranging between 70 and 87% sequence identity, according to high quality alignments) makes barcoding de-multiplexing a difficult task in nanopore data. However, our results indicate that with the configuration and parameters presented here we could efficiently distinguish the reads generated from HM782D and D6305 amplicons. As a consequence, the performance of this long amplicon approach to properly assign microbial communities to samples was efficiently assisted by the parameters during the de-multiplexing process that were central to discern reads obtained from respective samples multiplexed in the MinION flowcell. For instance, the distribution of reads matching with close related species such as *Lactobacillus gasseri* and *Lactobacillus fermentum*, contained distinctively in HM782D and D6305 samples, was indicative of the adequate execution of the de-multiplexing pipeline. The above was also exemplified for *Salmonella enterica* sequences that were determined only in D6305 despite its close relationship with *E. coli* at the 16S and 23S sequence level (close to 100%). Regarding the latter, the multiple sequence alignment built with *rrn* regions from both species was inspected directly distinguishing the ITS as the major source of variation between the two species. Indeed, this was corroborated by the comparative analysis performed during the clustering step of the reference samples to create our *rrn* database.

### Performance of R9.4 chemistry

During the course of the present work the MinION R9.4 chemistry was delivered in Autumn 2016. Therefore, we wanted to perform a replicate experiment using this type of chemistry in order to disclose how much improvement our approach would gain in terms of sensibility and specificity. With only 3h run we observed a notable improvement of throughput and per-base accuracy and the MinION™ produced almost 40,000 reads with a predominant QScore distribution between 8 and 12 suggesting a theoretical error rate of reads between 0.15 to 0.06, respectively, lower than obtained from R9 reads (0.25 to 0.15). After compiling all sequences in a fasta file, we proceeded to perform filtering in equal manner than previously done for R9 data. Consequently, we retained more than 33,000 reads (86%) for further processing and taxonomy assignment. The major results from comparison among R9 and R9.4 runs are summarized in the Table 2. As expected, the R9.4 dataset was more accurate and its reads showed a lower per-base error rate, therefore, the taxonomy analysis based on this reads would be more precise than observed with R9 reads. Globally, the results obtained from R9.4 chemistry are very similar than those observed with R9 chemistry but the level of uncertainty was diminished by reducing the number of close species to that contained in respective mock communities exhibiting very low abundance (<1%), thus decreasing from 40 species to 15 for the HM782D and from 18 to 16 for the D6305. We were unable again to recover *D. radiodurans* and *H. pylori* reads but we improved the sensitivity for *A. odontolyticus* and *R. sphaeroides* (Figure 3C and 3D), whose relative proportions were almost duplicated in R9.4 data (*R. sphaeroides* = 0.44%, *A. odontolyticus* = 0.31%). We compared the respective proportions obtained from R9 and R9.4 chemistries obtaining consistent results (Figure 3E) indicating that our approach is reproducible with no major changes despite the different chemistry and kits for library preparation using during both sequencing runs.

**Table 2.** Basic stats comparison of R9 and R9.4 reads after processing.

|  | **R9<sup>1</sup>** | **R9.4<sup>1</sup>** |
| --- | --- | --- |
| Total raw data | 17,038 (100%) | 39,216 (100%) |
| Reads > 1.5kb | 12,278 (72%) | 33,764 (86%) |
| Read length distribution | 25th percentile = 2,847nt<br>Median = 3,303nt<br>75th percentile = 3,754 | 25th percentile = 3,730nt<br>Median = 3,976nt<br>75th percentile = 4,135nt |
| Reads aligned (bc1 + bc8) | 3,227 (19%) | 14,392 (43%) |
| Alignment identity distribution | 25th percentile = 66.5%<br>Median = 69 %<br>75th percentile = 73% | 25th percentile = 81%<br>Median = 85%<br>75th percentile = 87% |
| Maximum identity | 86.7% | 92% |

1 The experiment and run accessions for the R9 data at ENA are ERX1676087 and ERR1605520, respectively.
2 The experiment and run accessions for R9.4 data at ENA are ERX1981944 and ERR1924217, respectively.

### Comparison with Illumina MiSeq data

The Illumina MiSeq data obtained after sequencing the V4-V5 16S region permitted to characterize the genus distribution in the HM782D sample with the RDP Classifier. As a result, we compiled distribution of all 17 genus represented in the HM782D mock community (Supplementary Material 1) and 4 additional genus with very low abundance (2 reads / 8,409 assigned). Moreover, when a OTU-picking approach was conducted we recovered 41 OTUs whose identity was evaluated in the SINA server (Supplementary Material 2). Globally, we recovered taxonomy identification of all genus expected but only three species were well identified based on the Greengenes taxonomy (*H. pylori, P. acnes*, and *R. sphaeroides*) whereas one was wrongly identified (*Neisseria cinerea*). For the D6305 mock community we could recover the eight different component at genus level of this mock community (Supplementary Material 1) plus seven additional and not related genus with very low abundance (< 7 reads / 8,046 assigned). At OTUs level, we retrieved a total of 14 sequences whose taxonomy identification is presented in the Supplementary Material 3. In this case, only *S. enterica* could be identified at species level. Given that data derived from this short read approach normally cannot reach a reliable taxonomy assignment down to species level, we proceed to make comparisons with R9 and R9.4 data by compiling these last information to genus level in order to evaluate the performance of our approach with a commonly used procedure. In the Supplementary Material 1 and Table 3 a comparison in terms of the relative read proportion and coverage bias is depicted. We observed no larger deviations in data retrieved with MinION regarding those numbers obtained with conventional approaches such as study of V4-V5 regions with MiSeq platform. Interestingly, we observed similar pattern of important negative coverage bias in all three approaches for *Actinomyces* spp., *Enterococcus* spp., and *Rhodobacter* spp. species in the HM782D community and for *Lactobacillus* spp., and *Listeria* spp., in the D6305 community, then suggesting that species of such genera are equally underrepresented no matter the type of amplicon, sequencing platform, or sequencing chemistry of study. Conversely, only the *Bacillus* spp. species from the D6305 exhibited large positive coverage bias values in all three approaches. Globally, all methods compared to study microbial communities at this level have a pattern of underrepresentation for all species present in the mock communities given the average and median values obtained. Moreover, the MiSeq V4-V5 approach also showed important coverage bias indicating this issue is not strictly associated with the MinION™ based approach presented in here and probably it is inherent to the amplification process of target DNA. Finally, correlation tests indicate that despite of coverage bias observed all configurations used to study the mock communities replicate fairly well the composition of the mock communities and that data obtained from R9 and R9.4 experiments show a slight improvement at this regard with no major differences when compared with data from MiSeq platform (Table 3).

**Table 3.** Comparative analysis among data generated from MinION and MiSeq platforms.

| Genera HM782D | Relative read proportion |  |  |  | Coverage bias |  |  |
| --- | --- | --- | --- | --- | --- | --- | --- |
|  | Reference | PCR+MiSeq | PCR+MinION-R9 | PCR+MinION-R9.4 | PCR+MiSeq | PCR+MinION-R9 | PCR+MinION-R9.4 |
| *Acinetobacter spp.* | 0.050 | 0.019 | 0.046 | 0.032 | -1.42 | -0.12 | -0.62 |
| *Actinomyces spp.* | 0.050 | 0.010 | 0.001 | 0.003 | -2.36 | -5.36 | -4.02 |
| *Bacillus spp.* | 0.050 | 0.017 | 0.102 | 0.045 | -1.57 | 1.03 | -0.16 |
| *Bacteroides spp.* | 0.050 | 0.106 | 0.059 | 0.037 | 1.08 | 0.25 | -0.42 |
| *Clostridium spp.* | 0.050 | 0.125 | 0.027 | 0.032 | 1.32 | -0.91 | -0.66 |
| *Deinococcus spp.* | 0.050 | 0.109 | 0.000 | 0.000 | 1.12 | ND | ND |
| *Enterococcus spp.* | 0.050 | 0.022 | 0.012 | 0.013 | -1.17 | -2.04 | -1.93 |
| *Escherichia/Shigella spp.* | 0.050 | 0.038 | 0.172 | 0.209 | -0.38 | 1.79 | 2.07 |
| *Helicobacter spp.* | 0.050 | 0.040 | 0.000 | 0.000 | -0.32 | ND | ND |
| *Lactobacillus spp.* | 0.050 | 0.065 | 0.051 | 0.068 | 0.38 | 0.03 | 0.45 |
| *Listeria spp.* | 0.050 | 0.024 | 0.074 | 0.133 | -1.04 | 0.57 | 1.41 |
| *Neisseria spp.* | 0.050 | 0.099 | 0.064 | 0.056 | 0.98 | 0.36 | 0.17 |
| *Propionibacterium spp.* | 0.050 | 0.021 | 0.079 | 0.097 | -1.25 | 0.66 | 0.96 |
| *Pseudomonas spp.* | 0.050 | 0.038 | 0.018 | 0.079 | -0.39 | -1.46 | 0.67 |
| *Rhodobacter spp.* | 0.050 | 0.013 | 0.002 | 0.004 | -1.91 | -4.36 | -3.51 |
| *Staphylococcus spp.* | 0.100 | 0.037 | 0.125 | 0.086 | -1.44 | 0.32 | -0.22 |
| *Streptococcus spp.* | 0.150 | 0.217 | 0.115 | 0.093 | 0.53 | -0.38 | -0.69 |
| **Genera D6305** |  |  |  |  |  |  |  |
| *Bacillus spp.* | 0.139 | 0.574 | 0.383 | 0.340 | 2.04 | 1.46 | 1.29 |
| *Enterococcus spp.* | 0.088 | 0.078 | 0.057 | 0.082 | -0.17 | -0.64 | -0.11 |
| *Escherichia/Shigella spp.* | 0.113 | 0.035 | 0.167 | 0.137 | -1.67 | 0.56 | 0.28 |
| *Lactobacillus spp.* | 0.198 | 0.104 | 0.049 | 0.049 | -0.93 | -2.01 | -2.01 |
| *Listeria spp.* | 0.163 | 0.046 | 0.080 | 0.118 | -1.82 | -1.03 | -0.46 |
| *Pseudomonas spp.* | 0.058 | 0.060 | 0.038 | 0.039 | 0.05 | -0.62 | -0.56 |
| *Salmonella spp.* | 0.115 | 0.049 | 0.099 | 0.138 | -1.22 | -0.22 | 0.26 |
| *Staphylococcus spp.* | 0.126 | 0.051 | 0.094 | 0.095 | -1.30 | -0.42 | -0.41 |
| **Average** | **0.080** | **0.080** | **0.077** | **0.079** | **-0.51** | **-0.55** | **-0.36** |
| **Median** | **0.050** | **0.048** | **0.062** | **0.074** | **-0.72** | **-0.30** | **-0.29** |
| **Min** | **0.050** | **0.010** | **0.000** | **0.000** | **-2.36** | **-5.36** | **-4.02** |
| **Max** | **0.198** | **0.574** | **0.383** | **0.340** | **2.04** | **1.79** | **2.07** |
| **Pearson's $r^a$ (p-value)** | -- | **0.39 (0.0504)** | **0.43 (0.0306)** | **0.41 (0.0417)** |  |  |  |
| **Pearson's $r^b$ (p-value)** | -- | -- | **0.73 (0.0001)** | **0.64 (0.0005)** |  |  |  |

a Pearson's r calculated from comparisons of R9, R9.4, and MiSeq data with reference proportions, respectively.
b Pearson's r calculated from comparisons of R9 and R9.4 data with MiSeq output, respectively.

## Discussion

The inventory of microbial species based on 16S rDNA sequencing is frequently used in biomedical research to determine microbial organisms inhabiting the human body and their relationship with disease. Recently, third-generation of DNA sequencing platforms have developed rapidly, facilitating the identification of microbial species and overcoming the read-length issues inherent to second-generation sequencing methods. These advances allow researchers to infer taxonomy and analyse diversity from the almost full-length bacterial 16S rRNA sequence [11, 14, 15, 17]. Particularly, the ONT platform deserves special attention given its portability and its fast development since the MinION™ became available in 2014. Notwithstanding, this technology is susceptible to a large stochastic variation, essentially derived from the wet-lab methods [1]. We corroborated this issue by obtaining a sequencing run where the raw data predominantly consisted of 1D reads as a consequence of the HP adapter ligation failure, despite following the manufacturer’s instructions. However, we were able to develop an efficient analysis protocol where the higher read quality offered by R9 chemistry and the updated Metrichor basecaller protocol proved pivotal to obtain 1D reads with a range of identity between 70 and 86%, with sufficient per-base accuracy to successfully perform the taxonomic analyses described herein. Moreover, during the course of this study the R9.4 flowcells were released and we were able to replicate our approach using this improved pore chemistry and the SQK-LSK108 for 1D libraries obtaining reads with sequence identity up to 92%.

Our preliminary results indicated that the *rrn* region in bacteria preferentially has a unique conformation (with the transcriptional arrangement of 16S-ITS-23S) and we could amplify this ~4.5Kbp region with the selected S-D-Bact-0008-c-S-20 and 23S-2241R primer pair. Once we were able to distinguish the feasibility to amplify the *rrn*, our approach comprised the study of two different mock communities in a multiplex manner, to be combined in one single MinION™ flowcell. By designing the respective forward and reverse primers tagged with specific barcodes recommended by ONT, we were able to retrieve extended barcode associated reads, in spite of the large proportion of per-base errors contained in these types of reads. Using MinION™ data based on multi-locus markers and long amplicon sequencing, we could reconstruct the structure of two commercially available mock communities. Although the expected proportions of some species in each community exhibited an important coverage bias, we were able to recover 80% (HM782D) and 100% (D6305) of bacterial species from the respective mock communities. Consequently, future analyses should be conducted to find an appropriate PCR approach using primers with a higher coverage for bacterial species.

We have analysed a great amount of genetic information with the aim of compiling a valuable database containing the genetic information for the *rrn* present in over 67,000 draft and complete bacterial genomes. The global length distributions in the region indicated that the *rrn* was 4,993 ± 187 bp in length whereas the 16S, ITS, and 23S subregions were 1,612 ± 75, 488 ± 186, and 3,036 ± 160 bp in length, respectively. Using this genetic information of the *rrn* and clustered at 100% of sequence identity enabled us to establish a multi-locus marker able to discriminate the taxonomy of two mock communities containing very close species. The latter was possible given that simultaneous analysis of the 16S, ITS, and 23S molecules offered almost 40-fold more diversity that studying the 16S, ITS, or 23S sequences separately and at 97% sequence identity. Moreover, the ITS was distinguished individually as an important variable genetic region in terms of sequence and length. Furthermore, it contributes notably to the higher variability observed in the *rrn* region, a fact evidenced in previous studies [18-21]. The accumulation of a larger number of variable sites in the *rrn* region, together with the particular structural variation of the ITS to potentially accommodate and encode tRNA genes, are thought to be central to discriminating bacterial species, despite the large proportion of per-base errors contained in MinION™ reads. Our data indicate that our MinION reads produce alignments with averaged length of 2,463 and 3,191 bases for HM782D and D6305, respectively, using R9 chemistry and 4,173 and 4,115 bases for HM782D and D6305, respectively, using R9.4 chemistry. Consequently, the taxonomy assignment was predominantly based on the variability of more than two out of the three markers included in the *rrn*, no matter if reads were produced from the 16S or 23S edges of *rrn* amplicons. We expect this type of analysis will likely become more accurate over time as nanopore chemistry improves in near future, with the concomitant increase in throughput, which is pivotal to disclose the hundreds of species present in complex microbial communities for analysis in human or environmental studies. Therefore, the multi-locus, long and multiplex methods described here represent a promising analysis routine for microbial and pathogen identification, relying on the sequence variation accumulated in approximately 5kbp of DNA, roughly accounting for the assessment of 1.25% of an average bacterial genome (~4Mbp). Notwithstanding, we cannot obviate that the current state of this approach presents some limitations in terms of the completeness of the *rrn* database created as well as the efficiency of the primers used to generate the long amplicons that have to be revisited in order to improve and increase the coverage of bacterial species. At date, our database include *rrn* sequences from 2,479 different species grouped into 918 different genus. In consequence, urgent studies must be undertaken to generate a more complete database including the *rrn* genomic information from species inhibiting complex and real samples such as those derived from human body.

## Methods

### Bacterial DNA and rrn amplicons

The complex DNA sample for preliminary studies of *rrn* region arrangement consisted of DNA isolated from faeces, kindly donated by a healthy volunteer upon informed consent. An aliquot of 200 mg of human faeces was used to isolate microbial DNA using the QIAamp DNA Stool Mini Kit (Qiagen) and following the manufacturer’s instructions. Finally, DNA was eluted in 100 μL nuclease-free water and a DNA aliquot at 20 ng/μL was prepared for PCR reaction using the primer pairs S-D-Bact-0008-c-S-20 and 23S-2241R or 23S-129F and S-D-Bact-1391-a-A-17 for testing configurations shown in Figure 1A (Table 1). The band size was analysed using the Java-based GelAnalyzer tool (www.gelanalyzer.com). Genomic DNA for the reference mock microbial communities was kindly donated by BEI Resources (http://www.beiresources.org) and ZYMO Research Corp (http://www.zymoresearch.com). The composition of the mock communities was as follows: i) HM782D is a genomic DNA mixture of 20 bacterial species containing equimolar ribosomal RNA operon counts (100,000 copies per organism per μL), as indicated by the manufacturer; and ii) ZymoBIOMICS Cat No. D6305 (D6305 hereinafter) is a genomic DNA mixture of eight bacterial species (and two fungal species) presented in equimolar amounts of DNA. According to manufacturers’ instructions, 1 μL of DNA from each mock community was used to amplify all the genes contained in the *rrn*. DNA was amplified in triplicate by 27 PCR cycles at 95°C for 30 s, 49°C for 15 s, and 72°C for 210 s. Phusion High-Fidelity Taq Polymerase (Thermo Scientific) and the primers S-D-Bact-0008-c-S-20 (mapping on 5′ of 16S gene) and 23S-2241R (mapping on 3′ of 23S gene), which target a wide range of bacterial 16S rRNA genes [22, 23]. For the Illumina MiSeq sequencing the V4-V5 hypervariable regions from bacterial 16S rRNA gene were amplified using 1 μL of DNA from each mock community and 25 PCR cycles at 95°C for 20 s, 40°C for 30 s, and 72°C for 20 s. Phusion High-Fidelity Taq Polymerase (Thermo Scientific) and the 6-mer barcoded primers S-D-Bact-0563-a-S-15 (AYTGGGYDTAAAGNG) and S-D-Bact-0907-a-A-20 (CCGTCAATTYMTTTRAGTTT). As we wished to multiplex the sequencing of both mock communities into one single MinION™ flowcell, we designed a dual-barcode approach where respective primers were synthesized and fused with two different barcodes recommended by ONT (Table 1). Amplicons consisted of ~4.5kbp blunt-end fragments for MinION approach and ~380bp for Illumina MiSeq approach, and those were purified using the Illustra GFX PCR DNA and Gel Band Purification Kit (GE Healthcare). Amplicon DNA was quantified using a Qubit 3.0 fluorometer (Life Technologies). Quantification of certain PCR products to correlate with sequencing coverage bias by qPCR was assessed according to previously described [11].

### Amplicon DNA library preparation

The Genomic DNA Sequencing Kit SQK-MAP006 was ordered from ONT and used to prepare the amplicon library for loading into the MinION™. Approximately 0.9 μg of amplicon DNA (0.3 per mock community plus 0.3 μg of an extra query sample consisting amplicons obtained from a genomic DNA mix of several uncharacterized microbial isolates) were processed for end repair using the NEBNextUltra II End Repair/dA-tailing Module (New England Biolabs), and followed by purification using Agencourt AMPure XP beads (Beckman Coulter) and washing twice with 1 volume of fresh 70% ethanol. Subsequently, and according to the manufacturer’s suggestions, we used 0.2 pmol of the purified amplicon DNA (~594 ng, assuming fragments of 4.5kbp in length) to perform the adapter ligation step. Ten μL of adapter mix, 2 μL of HP adapter, and 50 μl of Blunt/TA ligase master mix (New England Biolabs) were added in that order to the 38 μl end-repaired amplicon DNA. The reaction was incubated at room temperature for 15 minutes, 1 μL HP Tether was added and incubated for an additional 10 minutes at room temperature. The adapter-ligated amplicon was recovered using MyOne C1-beads (Life Technologies) and rinsed with washing buffer provided with the Genomic DNA Sequencing Kit SQK-MAP006 (Oxford Nanopore Technologies). Finally, the sample was eluted from the MyOne C1-beads by adding 25 μL of elution buffer and incubating for 10 minutes at 37°C before pelleting in a magnetic rack. The R9.4 sequencing library was obtained by processing of 600 ng of purified amplicon DNA (0.15 per mock community plus 0.15 μg of two extra query sample consisting amplicons obtained from a genomic DNA mix of several uncharacterized microbial isolates) with the SQK-LSK108 (Oxford Nanopore Technologies) sequencing for 1D reads and following the manufacturer’s instructions. Briefly, the 600 ng of amplicon DNA diluted in 50 μL nuclease-free water were processed for end repair using the NEBNextUltra II End Repair/dA-tailing Module (New England Biolabs), and followed by purification using Agencourt AMPure XP beads (Beckman Coulter) and washing twice with 200 μL volume of fresh 70% ethanol. The ligation step was performed with 30 μL of DNA end-prepped, 20 μL adapter mix, and 50 μl of Blunt/TA ligase master mix (New England Biolabs). The reaction was incubated at room temperature for 15 minutes at room temperature. The adapter-ligated amplicon was recovered again Agencourt AMPure XP beads (Beckman Coulter), washing twice with the ABB buffer supplied into the SQK-LSK1008 sequencing kit (Oxford Nanopore Technologies), and eluted from Agencourt AMPure XP beads by adding 25 μL of elution buffer and incubating for 10 minutes at 37°C before pelleting in a magnetic rack. Samples for Illumina MiSeq approach were sent to the National Center for Genomic Analaysis (CNAG, Spain) for multiplex sequencing in one lane of MiSeq instrument with 2x300 paired-end configuration.

### Flowcell set-up

A brand new, sealed R9 flowcell was acquired from ONT and stored at 4°C before use. The flowcell was fitted to the MinION™ MkIb prior to loading the sequencing mix, ensuring good thermal contact. The R9 flowcell was primed twice using 71 μL premixed nuclease-free water, 75 μL 2x running buffer, and 4 μL fuel mix. At least 10 minutes were required to equilibrate the flowcell before each round of priming and before final DNA library loading. For the replicate experiment, a R9.4 flowcell was fitted to the MinION™ MkIb prior to loading the sequencing mix, ensuring good thermal contact. The R9.4 flowcell was primed with 800 μL of running buffer (0.5 mL nuclerase-free water plus 0.5 mL RBF buffer). At least 10 minutes were required to equilibrate the flowcell and then the remaining 200 μL of running buffer were injected into the R9.4 flowcell with the SpotON port opened.

### Amplicon DNA sequencing

The sequencing mix was prepared with 59 μl nuclease-free water, 75 μl 2x running buffer, 12 μL DNA library, and 4 μL fuel mix. A standard 48-hour sequencing protocol was initiated using the MinKNOW™ v0.51.3.40. Base-calling was performed through data transference using the MetrichorTM agent v2.40.17 and the Barcoding plus 2D Basecalling RNN for SQK-NSK007 workflow v1.107. During the sequencing run, one additional freshly diluted aliquot of DNA library was loaded after 16 hours of initial input. The raw sequencing data derived from the two mock communities studied here was expected to account two-thirds of the data produced by the R9 flowcell used. The R9.4 run was performed with 75 μL DNA library (37.5 μL, RBF buffer, 25.5 μL LLB, 12 μL DNA library) loaded into the R9.4 flowcell through the SpotON port. A standard 48-hour sequencing protocol was initiated using the MinKNOW™ v1.5.5 with the respective local basecalling algorithm implemented in the MinKNOW™ software. The R9.4 raw data was generated during only 3h sequencing run.

### The rrn database

We built a database containing the genetic information for the 16S and 23S rRNA genes and the ITS sequence in all the complete and draft bacterial genomes available in the NCBI database (ftp://ftp.ncbi.nlm.nih.gov/genomes/genbank/bacteria). A total of 67,199 genomes were analysed by downloading the “*fna*” files and parsing for rRNA genes into the respective “*gff*” annotation file. Chromosome coordinates for *rrn* regions were parsed and used to extract such a DNA sequences from complete chromosomes or DNA contigs assembled. The resulting *rrn* sequences were analysed and the length distribution was assessed. We retrieved a total of 47,698 *rrn* sequences with an average of 4,993 nt in length. By selecting the size distribution equal to the 99th percentile (two-sided), we discarded potential incomplete or aberrant annotated *rrn* sequences and observed that *rrn* sequences can be found between 4,196 and 5,790nt; under these boundaries, our *rrn* database finally accounted for a total of 46,920 sequences. Equivalent databases were built by parsing the respective *rrn* sequences with the RNammer tool to discriminate the 16S, ITS, and 23 S rRNA sequences [24]. To remove the level of redundancy of our *rrn* database and to maintain the potential discriminatory power at strain level, we performed clustering analysis using USEARCH v8 tool for sequence analysis and the option *-otu_radius_pct* equal 0 [25], thus obtaining a total of 22,350 reference sequences. For comparative aims, the *rrn* database and the 16S, ITS, and 23S databases were also analysed using the option *-otu_radius_pct* with values ranging from 1 to 3. For accessing to *rrn* database and the respective species annotation, see Availability of supporting data.

### MinION data analysis

Read-mapping was performed using the LAST aligner v.189 [26] with parameters -q1 -b1 -Q0 -a1 -r1. Each 1D read was compared in a competitive way against the entire *rrn* database and the best hit was selected by obtaining the highest alignment score. Alignment length as well as alignment coordinates in target and query sequences were parsed from the LAST output and the sequence identity between matched regions was calculated using the python *Levenshtein* distance package. An iterative processing was used to determine thresholds for detection by evaluating the taxonomy distribution with reads subsampling and different levels of sequence identity in top scored alignments. High quality alignments were selected by filtering out those with identity values up to the 50th percentile of the distribution of identity values of all reads per sample (~69%) in the R9 run. Therefore, taxonomy assignment was based exclusively on alignments with ≥ 70% identity. For data derived from R9.4 chemistry, high quality alignments were selected by filtering out those with identity values up to 25th percentile of the distribution, thus retaining alignments with ≥ 81% identity. Basic stats, distributions, filtering, and comparisons were performed in R v3.2.0 (https://cran.r-project.org). For relative quantification of species the singletons were removed and the microbial species considered to be predominantly present in the mock communities were those with a relative a proportion ≥ 1%, a value that demonstrated to be discriminative to always obtain the expected microbial diversity during the iterative processing of alignments. The coverage bias was calculated by obtaining fold-change (Log_2_) of species-specific read counting against the expected (theoretical) average for the entire community according to information provided by the manufacturers.

### Illumina data analysis

Fastq paired-end raw data (ENA experiment accession ERX2062322) was were assembled using *Flash* software [27]. The HM782D and D6305 reads were de-multiplexed and barcode and primers were removed using *Mothur* v1.36.1 [28]. The sequences were then processed for chimera removal using *Uchime* algorithm [29] and SILVA reference set of 16S sequences [30]. A normalized subset of 10,000 sequences per sample was created by random selection after shuffling (10,000X) of the original dataset. Taxonomy assessment was performed using the RDP classifier v2.7 [31]. The Operational Taxonomic Unit (OTU)-picking approach was performed with the normalized subset of 10,000 sequences and the *uclust* algorithm implemented in USEARCH v8.0.1623 and the options *-otu_radius_pct* equal 3 for clustering at 97% and *-min*size 2 for remove singletons [25]. SINA server was used for taxonomy identification of OTUs recovered from Illumina MiSeq data [32].

## Availability of supporting data

Accessions for the *rrn* database containing the reference sequences for alignments and taxonomic annotation is available at https://github.com/alfbenpa/rrn_db. The code source of the original *split_barcodes.pl* perl script is available at https://github.com/nanoporetech/barcoding/releases/tag/1.0.0 with ONT copyright.

## Abbreviations

EC: European Commission
ENA: European Nucleotide Archive
HDF: Hierarchical Data Format
ITS: internal transcribed spacer
NCBI: National Center for Biotechnology Information
ONT: Oxford Nanopore Technologies
PCR: Polymerase Chain Reaction
rDNA: DNA encoding for the Ribosomal RNA
rRNA: Ribosomal RNA
*rrn*: the DNA region containing the 16S and 23S bacterial rRNA genes and its respective ITS region
SINA: SILVA Incremental Aligner
USB: Universal Serial Bus

## Competing interests

ABP is part of the MinION™ Access Programme (MAP).

## Authors’ contributions

ABP and YS designed the study and managed the project. ABP performed the experiments, analysed and managed the data. ABP draft the manuscript. Both authors read and approved the final manuscript.

## Acknowledgements

This work and the contract to ABP is supported by the European Union’s Seventh Framework Program under the grant agreement n° 613979 (MyNewGut).

**Supplementary Material 1.** Comparison of MinION™ and MiSeq outputs. Data obtained from Illumina MiSeq sequencing of V4-V5 16S region from respective mock communities was compared with outputs from MinION™ R9 and R9.4. Given that taxonomy identification of MiSeq reads at species level only retrieved very few assignments, we compiled the species distribution of MinION™ data at genus level.

**Supplementary Material 2.** MS Excel file compiling the output information retrieved from SINA server (https://www.arb-silva.de/aligner/) for taxonomy identification of 41 OTUs derived from analysis of HM782D with Illumina MiSeq approach. Information regarding sequence quality, identity percentage, mapping coordinates against *E. coli* reference, length and taxonomy based on SILVA, Greengenes, and RDP databases is showed for all OTU.

**Supplementary Material 3.** MS Excel file compiling the output information retrieved from SINA server (https://www.arb-silva.de/aligner/) for taxonomy identification of 18 OTUs derived from analysis of D6305 with Illumina MiSeq approach. Information regarding sequence quality, identity percentage, mapping coordinates against *E. coli* reference, length and taxonomy based on SILVA, Greengenes, and RDP databases is showed for all OTU.

## References

1. Ip CL, Loose M, Tyson JR, de Cesare M, Brown BL, Jain M, Leggett RM, Eccles DA, Zalunin V, Urban JM et al: MinION Analysis and Reference Consortium: Phase 1 data release and analysis. F1000Res 2015, 4:1075.

2. Loman NJ, Quick J, Simpson JT: A complete bacterial genome assembled de novo using only nanopore sequencing data. Nat Methods 2015, 12(8):733–735.

3. Karlsson E, Larkeryd A, Sjodin A, Forsman M, Stenberg P: Scaffolding of a bacterial genome using MinION nanopore sequencing. Sci Rep 2015, 5:11996.

4. Risse J, Thomson M, Patrick S, Blakely G, Koutsovoulos G, Blaxter M, Watson M: A single chromosome assembly of Bacteroides fragilis strain BE1 from Illumina and MinION nanopore sequencing data. Gigascience 2015, 4:60.

5. Ammar R, Paton TA, Torti D, Shlien A, Bader GD: Long read nanopore sequencing for detection of HLA and CYP2D6 variants and haplotypes. F1000Res 2015, 4:17.

6. Norris AL, Workman RE, Fan Y, Eshleman JR, Timp W: Nanopore sequencing detects structural variants in cancer. Cancer Biol Ther 2016:1–8.

7. Greninger AL, Naccache SN, Federman S, Yu G, Mbala P, Bres V, Stryke D, Bouquet J, Somasekar S, Linnen JM et al: Rapid metagenomic identification of viral pathogens in clinical samples by real-time nanopore sequencing analysis. Genome Med 2015, 7(1):99.

8. Kilianski A, Haas JL, Corriveau EJ, Liem AT, Willis KL, Kadavy DR, Rosenzweig CN, Minot SS: Bacterial and viral identification and differentiation by amplicon sequencing on the MinION nanopore sequencer. Gigascience 2015, 4:12.

9. Ashton PM, Nair S, Dallman T, Rubino S, Rabsch W, Mwaigwisya S, Wain J, O’Grady J: MinION nanopore sequencing identifies the position and structure of a bacterial antibiotic resistance island. Nat Biotechnol 2015, 33(3):296–300.

10. Judge K, Harris SR, Reuter S, Parkhill J, Peacock SJ: Early insights into the potential of the Oxford Nanopore MinION for the detection of antimicrobial resistance genes. J Antimicrob Chemother 2015, 70(10):2775–2778.

11. Benitez-Paez A, Portune KJ, Sanz Y: Species-level resolution of 16S rRNA gene amplicons sequenced through the MinION portable nanopore sequencer. Gigascience 2016, 5:4.

12. Li C, Cheng KR, Boey JHE, Ng HQA, Wilm A, Nagarajan N: INC-Seq: Accurate single molecule reads using nanopore sequencing. bioRxiv 2016:doi: http://dx.doi.org/10.1101/038042

13. Myer PR, Kim M, Freetly HC, Smith TP: Evaluation of 16S rRNA amplicon sequencing using two next-generation sequencing technologies for phylogenetic analysis of the rumen bacterial community in steers. J Microbiol Methods 2016, 127:132–140.

14. Schloss PD, Jenior ML, Koumpouras CC, Westcott SL, Highlander SK: Sequencing 16S rRNA gene fragments using the PacBio SMRT DNA sequencing system. PeerJ 2016, 4:e1869.

15. Shin J, Lee S, Go MJ, Lee SY, Kim SC, Lee CH, Cho BK: Analysis of the mouse gut microbiome using full-length 16S rRNA amplicon sequencing. Sci Rep 2016, 6:29681.

16. Watson M, Thomson M, Risse J, Talbot R, Santoyo-Lopez J, Gharbi K, Blaxter M: poRe: an R package for the visualization and analysis of nanopore sequencing data. Bioinformatics 2014, 31(1):114–115.

17. Li C, Chng KR, Boey EJ, Ng AH, Wilm A, Nagarajan N: INC-Seq: accurate single molecule reads using nanopore sequencing. Gigascience 2016, 5(1):34.

18. Fernandez J, Avendano-Herrera R: Analysis of 16S-23S rRNA gene internal transcribed spacer of Vibrio anguillarum and Vibrio ordalii strains isolated from fish. FEMS Microbiol Lett 2009, 299(2):184–192.

19. Maslunka C, Gurtler V, Seviour R: Unusual features of the sequences of copies of the 16S-23S rRNA internal transcribed spacer regions of Acinetobacter bereziniae, Acinetobacter guillouiae and Acinetobacter baylyi arise from horizontal gene transfer events. Microbiology 2015, 161(Pt 2):322–329.

20. Stewart FJ, Cavanaugh CM: Intragenomic variation and evolution of the internal transcribed spacer of the rRNA operon in bacteria. J Mol Evol 2007, 65(1):44–67.

21. Tambong JT, Xu R, Bromfield ES: Intercistronic heterogeneity of the 16S-23S rRNA spacer region among Pseudomonas strains isolated from subterranean seeds of hog peanut (Amphicarpa bracteata). Microbiology 2009, 155(Pt 8):2630–2640.

22. Hunt DE, Klepac-Ceraj V, Acinas SG, Gautier C, Bertilsson S, Polz MF: Evaluation of 23S rRNA PCR primers for use in phylogenetic studies of bacterial diversity. Appl Environ Microbiol 2006, 72(3):2221–2225.

23. Klindworth A, Pruesse E, Schweer T, Peplies J, Quast C, Horn M, Glockner FO: Evaluation of general 16S ribosomal RNA gene PCR primers for classical and next-generation sequencing-based diversity studies. Nucleic Acids Res 2012, 41(1):e1.

24. Lagesen K, Hallin P, Rodland EA, Staerfeldt HH, Rognes T, Ussery DW: RNAmmer: consistent and rapid annotation of ribosomal RNA genes. Nucleic Acids Res 2007, 35(9):3100–3108.

25. Edgar RC: Search and clustering orders of magnitude faster than BLAST. Bioinformatics 2010, 26(19):2460–2461.

26. Frith MC, Hamada M, Horton P: Parameters for accurate genome alignment. BMC Bioinformatics 2010, 11:80.

27. Magoc T, Salzberg SL: FLASH: fast length adjustment of short reads to improve genome assemblies. Bioinformatics 2011, 27(21):2957–2963.

28. Schloss PD, Westcott SL, Ryabin T, Hall JR, Hartmann M, Hollister EB, Lesniewski RA, Oakley BB, Parks DH, Robinson CJ et al: Introducing mothur: open-source, platform-independent, community-supported software for describing and comparing microbial communities. Appl Environ Microbiol 2009, 75(23):7537–7541.

29. Edgar RC, Haas BJ, Clemente JC, Quince C, Knight R: UCHIME improves sensitivity and speed of chimera detection. Bioinformatics 2011, 27(16):2194–2200.

30. Quast C, Pruesse E, Yilmaz P, Gerken J, Schweer T, Yarza P, Peplies J, Glockner FO: The SILVA ribosomal RNA gene database project: improved data processing and web-based tools. Nucleic Acids Res 2013, 41(Database issue):D590–596.

31. Wang Q, Garrity GM, Tiedje JM, Cole JR: Naive Bayesian classifier for rapid assignment of rRNA sequences into the new bacterial taxonomy. Appl Environ Microbiol 2007, 73(16):5261–5267.

32. Pruesse E, Peplies J, Glockner FO: SINA: accurate high-throughput multiple sequence alignment of ribosomal RNA genes. Bioinformatics 2012, 28(14):1823–1829.

